# ProGeM: A framework for the prioritisation of candidate causal genes at molecular quantitative trait loci

**DOI:** 10.1101/230094

**Authors:** David Stacey, Eric B. Fauman, Daniel Ziemek, Benjamin B. Sun, Eric L. Harshfield, Angela M. Wood, Adam S. Butterworth, Karsten Suhre, Dirk S. Paul

## Abstract

Quantitative trait locus (QTL) mapping of molecular phenotypes such as metabolites, lipids, and proteins through genome-wide association studies (GWAS) represents a powerful means of highlighting molecular mechanisms relevant to human diseases. However, a major challenge of this approach is to identify the causal gene(s) at the observed QTLs. Here we present a framework for the “**Pr**ioritisation **o**f candidate causal **Ge**nes at **M**olecular QTLs” (ProGeM), which incorporates biological domain-specific annotation data alongside genome annotation data from multiple repositories. We assessed the performance of ProGeM using a reference set of 227 previously reported and extensively curated metabolite QTLs. For 98% of these loci, the expert-curated gene was one of the candidate causal genes prioritised by ProGeM. Benchmarking analyses revealed that 69% of the causal candidates were nearest to the sentinel variant at the investigated molecular QTLs, indicating that genomic proximity is the most reliable indicator of “true positive” causal genes. In contrast, *cis*-gene expression QTL data led to three false positive candidate causal gene assignments for every one true positive assignment. We provide evidence that these conclusions also apply to other molecular phenotypes, suggesting that ProGeM is a powerful and versatile tool for annotating molecular QTLs. ProGeM is freely available via GitHub.

## INTRODUCTION

With the continued application of genome-wide association studies (GWAS) to human disease aetiology (1-4), the rapid discovery rate of susceptibility loci is far outstripping the rate at which we are able to elucidate the biological mechanisms underlying the identified loci. This represents a major bottleneck to translational progress. Quantitative trait locus (QTL) mapping of molecular, intermediate phenotypes provides a powerful means to functionally annotate and characterise GWAS signals for complex traits in a high-throughput manner. This approach has been pioneered with the use of transcriptomic data to identify gene expression QTLs (eQTLs) (5-9). Recent technological advances have enabled the application of this approach to methylomic (10,11), proteomic (12-14), lipidomic (15), and metabolomic (16-18) data. This catalogue of molecular QTLs, cutting across multiple ‘omic modalities, can be readily queried to elucidate the functional impact of disease-associated variants not only on the abundance of transcripts, but also epigenetic marks, proteins, lipids, and metabolites.

A key challenge with these data relates to the identification of specific causal genes at the observed molecular QTLs. Accurate molecular QTL-gene assignments are critical for the meaningful interpretation of the biology underlying GWAS signals and the subsequent design of appropriate experimental follow-up work. There are several web tools available that facilitate the identification of genes most likely to be impacted functionally by either the sentinel or proxy variants tagging a molecular QTL. For example, tools such as the Single Nucleotide Polymorphisms Annotator (SNiPA) and the Functional Mapping and Annotation of GWAS tool (FUMA) integrate various positional, regulatory, and *cis*-eQTL datasets, enabling the identification of candidate causal genes using a data-driven approach (19,20). Nevertheless, the sensitivity of these tools, or the extent to which they are able to highlight “true positive” causal genes, has not yet been rigorously assessed. This is largely due to the time-consuming and resource-intensive experimental follow-up that is required to assign candidate causal genes at each individual association locus, resulting in a limited number of trait- or disease-associated variants that have so far been unequivocally assigned to established causal genes. Thus, “reference” causal gene sets for a particular trait of interest, which could be used to benchmark causal gene prioritisation tools, are either unavailable or likely to be unrepresentative.

However, metabolite QTL (mQTL) data represent a unique case in that there is an abundance of published biochemical mechanistic research pre-dating findings from GWAS that has identified and characterised many of the enzymes, transporters, and other proteins that regulate specific metabolites (21-23). In addition, this experimental research has been complemented by the study of numerous rare inborn errors of metabolism, thereby further elucidating the substrates and functions of metabolic gene products (24-26). Thus, by cross-referencing mQTLs identified by GWAS with this pre-existing body of biochemical and genetic research, it becomes possible to generate a large, high-confidence, mQTL-specific reference causal gene set, which can be used for the validation of both current and future causal gene prioritisation tools.

Another major limitation of these tools is that they are hampered by low specificity, in that they will typically highlight several candidate causal genes at most association loci. This necessitates the application of further downstream prioritisation methods or literature review. Therefore, causal gene prioritisation tools that are able to facilitate this process in an automated fashion will prove instrumental in refining the overall search space. As an example, the vast majority of current tools are not geared towards any specific trait or phenotype, relying solely on positional, regulatory, and/or *cis*-eQTL data to prioritise causal genes based on the likelihood that they are functionally affected by polymorphisms at a QTL or GWAS locus. Conversely, by designing a tool that is focused on a specific trait (e.g., metabolites) or trait class (e.g., molecular intermediates), relevant annotation data from publicly available databases (e.g., KEGG, GO) can be leveraged to directly prioritise those candidates that have been shown to regulate metabolites or other biomolecules.

Here we present an analysis framework and accompanying R script (https://github.com/ds763/ProGeM) for the **Pr**ioritisation **o**f candidate causal **Ge**nes at **M**olecular QTLs (ProGeM). Consistent with existing tools, ProGeM leverages positional and *cis*-eQTL data to prioritise genes most likely to be impacted functionally by variants tagging the molecular QTL. In addition, ProGeM integrates information from biological domain-specific annotation data from multiple repositories to prioritise genes involved in the biological mechanisms that regulate the molecular phenotype in question. In this way, ProGeM is able to harness both literature- and experimental-derived information in a quick and efficient manner. Crucially, and in contrast to existing tools, we have also determined the sensitivity and specificity of ProGeM using two molecular QTL datasets, comprising 227 mQTLs and 562 *cis*-pQTLs, for which each QTL has been assigned a high-confidence causal gene. Informed by these datasets, we make recommendations as to which annotation criteria may be most informative for the identification of candidate causal genes at molecular QTLs.

## MATERIAL AND METHODS

### Reference causal gene sets for molecular QTL data

*mQTL dataset*. Between 2007 and 2016, 109 papers reported results of a GWAS of metabolite levels. Suitable traits were identified largely through a manual review of all entries from the GWAS catalogue tagged with the Experimental Factor Ontology (EFO) term “measurement” (EFO_0001444) or any descendants of the term. This analysis focused on small molecules, ions, metabolites, vitamins, and other biomolecules not directly encoded by genes such as mRNA or proteins. The source tissue was most often plasma or serum although studies of urine and cerebrospinal fluid have also been included. Where available, full summary statistics for the identified studies were downloaded and peak-pruned to identify sentinel SNPs at least 1mb apart. This was done for datasets generated by the Global Lipids Genetics Consortium (i.e., LDL cholesterol, HDL cholesterol, total cholesterol, and total triglycerides), Meta-Analyses of Glucose and Insulin-related traits Consortium (i.e., fasting glucose), and TwinsUK/KORA (17), with a total of 78 metabolites with *p*≤5×10^−8^. Before clustering, there were 2,808 sentinel SNPs (*p*≤5×10^−8^) covering 250 distinct metabolites from these 109 studies. These variants were clustered into 497 loci by collapsing variants closer than 500 kilobases (kb) unless there was a compelling biochemical reason to separate the associated metabolites. For example, nine sentinel SNPs for branched chain amino acids and related metabolites near *PPM1K* were clustered together, and 21 sentinel SNPs for urate, uric acid, and urea near *ABCG2* were clustered. However, these two groups were not clustered further, even though they are only 150kb apart because the metabolites are not tightly linked biochemically and each cluster has its own very credible causal gene. Within each cluster, the variant with the smallest *p*-value (across all related metabolites) was retained.

For each of the 497 locus-metabolite pairs, all protein-coding genes within 1mb of the sentinel variant were considered. This generated a median of 20 genes per locus (range 4 to 92). While the final selection of the likely causal gene was performed manually following an expert review of the literature, a variety of approaches was employed to guide the literature review. For example, text-mining was used to identify papers mentioning the gene, gene product or synonyms, and metabolite or respective synonyms. The KEGG (Kyoto Encyclopedia of Genes and Genomes) pathway databases were mined for networks linking each gene at a locus to the associated metabolite. Many metabolites are so distinct that only a handful of genes in the whole genome have ever been discussed in relation to them. Examples include 5-oxoproline, here linked to the *OPLAH* gene that encodes 5-oxoprolinase (27), and manganese, here linked to the *SLC30A10* gene that encodes a manganese transporter (28).

For each of the 497 gene-metabolite pairs, we attempted to identify the earliest publication conclusively linking the gene product to the exact metabolite reported or a biochemically-similar metabolite. Preference was given to evidence for the human gene, to non-genetic data, and to experimental work conducted before the publication of the GWAS. The publications reporting the experimental validation for the causal genes are presented in **Supplementary Table S1**, listed under “Evidence Source (PMID)”.

*cis-pQTL dataset*. This dataset was derived from our recent large-scale pQTL study (14). Briefly, the dataset consisted of 3,301 healthy individuals of European descent, who had been randomly selected from a pool of ~50,000 participants of the INTERVAL study (29). Plasma protein levels were measured using the SOMAscan platform (SomaLogic, Inc., Boulder, Colorado, US) comprising 4,034 distinct aptamers (SOMAmers) covering 3,623 proteins (or protein complexes). Genotyping was performed using the Affymetrix Axiom UK Biobank genotyping array (Santa Clara, California, US), assaying ~830,000 variants. Variants were imputed using a combined 1000 Genomes Phase 3-UK10K reference panel, which yielded a total of ~10.5 million variants for pQTL analyses after stringent QC filtering. Overall, we found a total of 1,927 significant (*p*<1.5×10^−11^) genetic variant-protein associations involving 1,478 proteins and 764 unique genomic loci (14). Of these 1,927 associations, 555 were *cis*-associations (i.e., sentinel variant within 1mb of the gene encoding the corresponding protein) and the remaining 1,373 were *trans*-associations.

In order to benchmark our candidate causal gene identification strategy, we utilised only the *cis*-pQTL data, for which we hypothesised that the causal gene at a given *cis*-pQTL ought to be the gene that encodes the associated protein. To convert the 555 *cis*-pQTLs into a high-confidence set of sentinel variant-causal gene assignments, we first decomposed the *cis*-pQTLs into 589 sentinel variant-SOMAmer *cis*-associations. We then removed nine sentinel variants with associations originating from SOMAmers known to target more than a single protein due to paralogous sequences. We also removed 16 associations with SOMAmers that led to duplicate (or more) protein associations for the same sentinel variant. Finally, we removed two additional associations for which the same sentinel variants were associated with distinct protein isoforms encoded by single genes. Thus, we used a set of 562 high-confidence sentinel variant-causal gene assignments (**Supplementary Table S2**) for the purposes of benchmarking the bottom-up component of ProGeM.

### Proxy variant selection

We selected proxies for each sentinel variant based on an LD threshold of *r*^2^≥0.8. For the mQTL dataset, proxies were extracted from the 1000 Genomes Project (EUR Super Population) data using PhenoScanner, which is a curated database of publicly available results from large-scale genetic association studies (30). For the *cis*-pQTL dataset, proxies were derived directly from the genotype data of the participants, as previously described (14).

### Annotation of sentinel and proxy variants

All sentinel and proxy variants were annotated using the Ensembl Variant Effect Predictor (VEP) (v83) on GENCODE transcripts (v19) for GRCh37 (31). Annotations were generated using the “per gene” option, which considers variant annotations across all genes and transcripts, and selects the most severe consequence per gene with an arbitrary selection of the corresponding transcript. In particular, we made use of the IMPACT rating provided by VEP, which assigns input variants to one of four overarching functional categories as follows: (i) high impact: “the variant is assumed to have high (disruptive) impact on the protein, probably causing protein truncation, loss of function, or triggering nonsense mediated decay” (i.e., frameshift variant, start-lost variant); (ii) moderate impact: “a non-disruptive variant that might change protein effectiveness” (i.e., missense variant, inframe deletion); (iii) low impact: “assumed to be mostly harmless or unlikely to change protein behaviour” (i.e., synonymous variant, 3’-untranslated region variant); and (iv) modifier impact: “usually non-coding variants or variants affecting non-coding genes, where predictions are difficult or there is no evidence of impact” (i.e., intergenic variant, intronic variant).

### Identification of candidate causal genes

#### Bottom-up component

We used the GenomicRanges suite of R packages (32) to extract (i) the three nearest protein-coding genes to each sentinel variant, and (ii) any LD range-overlapping genes from a GRCh37 gene model based on a GTF file (“Homo_sapiens.GRCh37.82.gtf”) retrieved from Ensembl (33). LD ranges for each sentinel variant were defined as the range between the genomic coordinates (GRCh37) of the left- and right-most proxy variants (+/-5kb). In cases where the sentinel had no proxies, the coordinates of the sentinel variant (+/-5kb) were taken as the LD range. We also extracted significant *cis*-eQTL target genes of either sentinel or proxy variants from the *cis*-eQTL data prepared by the Genotype-Tissue Expression (GTEx) project (5) (v7), across all tissues assayed (*n*=48). Significant *cis*-eQTLs were defined by beta distribution-adjusted empirical *p*-values using a false discovery rate (FDR) threshold of 0.05 (see http://www.gtexportal.org/home/documentationPage for details).

#### Top-down component

mQTL sentinel variant-flanking genes (i.e., transcription start site (TSS) within +/-500kb of a sentinel) were identified using GenomicRanges (32) and the same Ensembl GTF file as above. Top-down candidates were then identified by cross-referencing the resultant list of sentinel-flanking genes against a list of known metabolic-related genes derived from five open-source databases (**Supplementary Table S3**).

### Statistical analyses

#### Sensitivity and specificity

Sensitivity was calculated and expressed as a percentage of the total number of molecular QTLs in question, as follows:

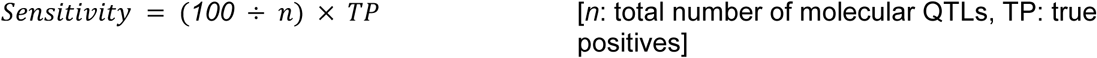

We calculated the overall specificity of ProGeM as:

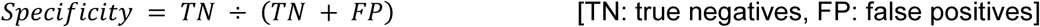

whereby maximal specificity is indicated by specificity = 1 and where TN was comprised of:

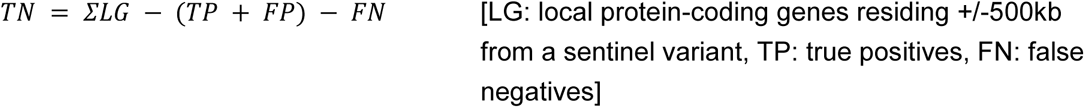

To compare the specificity of the bottom-up and top-down components, as well as the concurrent candidate gene sets, TN was comprised of:

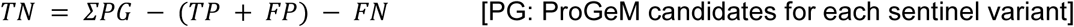

#### Enrichment analyses

Enrichment analyses were performed using Fisher’s exact tests, with the relevant background gene sets consisting of all remaining candidate causal genes (both bottom-up and top-down) across either the mQTL or *cis*-pQTL dataset as appropriate.

### Software for analyses

All analyses described in this study were performed using R v3.3.2 and Bioconductor v3.3.

## RESULTS

### Conceptual framework of ProGeM

We devised a general framework for the **Pr**ioritisation **o**f candidate causal **Ge**nes at **M**olecular QTLs (ProGeM) (**Figure 1**). This framework is based on the assumption that in order for a gene to be causal for a molecular QTL, or indeed any other phenotype, it must fulfil two requirements: (i) the gene product must exhibit altered structure, abundance or function as a result of the sentinel or proxy variants at the QTL, and (ii) the gene must be involved in the molecular mechanism that influences the trait in question. Accordingly, ProGeM is comprised of a “bottom-up” and a “top-down” component that prioritises candidate causal genes from the perspective of the genetic variant and the molecular phenotype, respectively.

**Figure 1.**
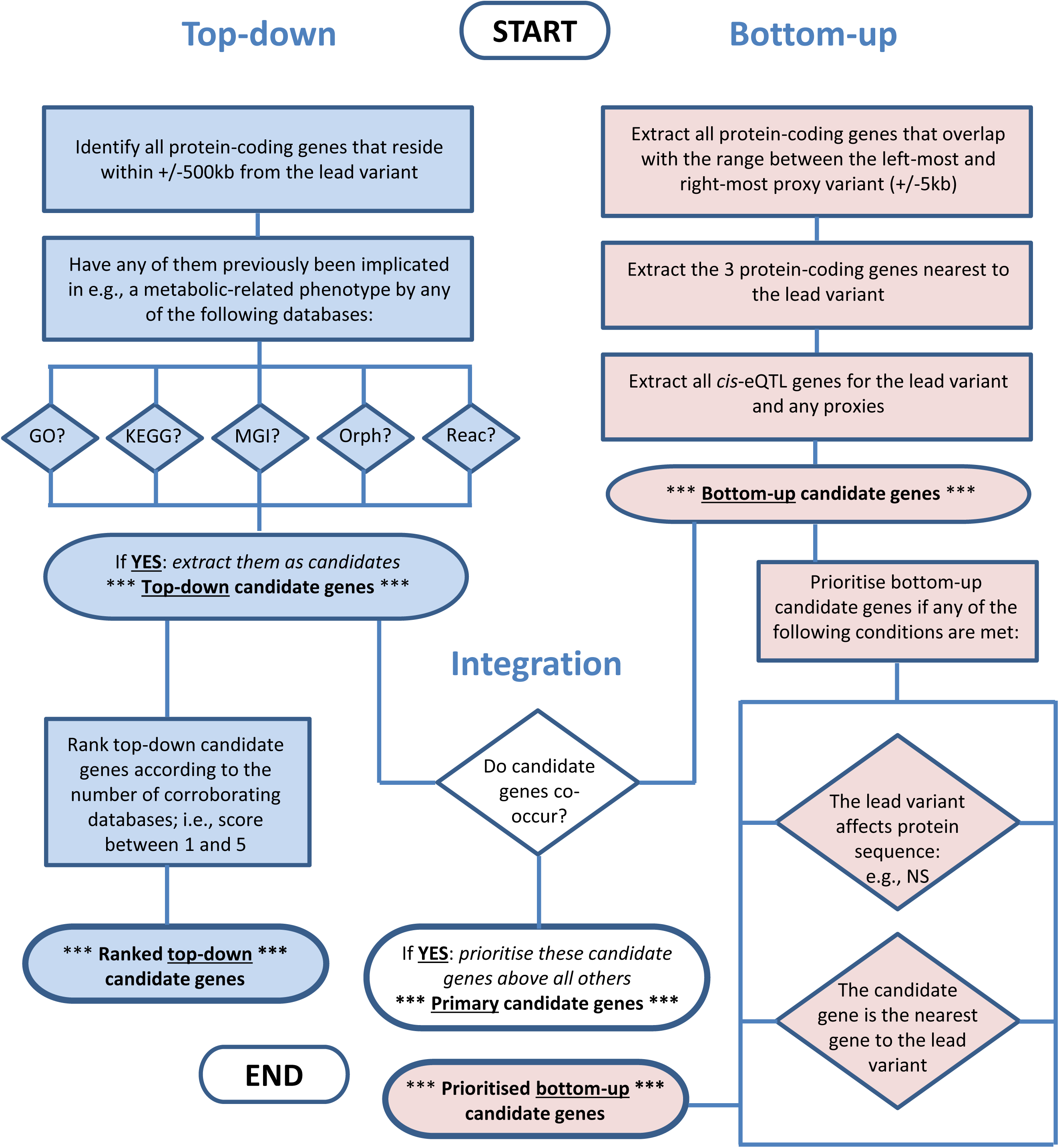
ProGeM: a framework for identifying and prioritising candidate causal genes at molecular QTL. A proxy is defined as those variants with *r*^2^≥0.8 with the sentinel variant (1000 Genomes Project, EUR Super Population). GTEx v7 data were used as a source for identifying *cis*-eQTLs. GO; Gene Ontology, KEGG; Kyoto Encyclopedia of Genes and Genomes, MGI; Mouse Genome Informatics, Orph; Orphanet, Reac; Reactome, NS; nonsynonymous.

#### Bottom-up component

For the bottom-up component, we utilise three complementary methods to identify plausible candidate causal genes based on (i) their proximity to the LD range (**Material and Methods**); (ii) their proximity to the sentinel variant; and (iii) whether their mRNA expression levels are impacted by either the GWAS sentinel or any corresponding proxy variants (**Figure 1**). The former two methods are designed to capture candidate causal genes that are proximal to the association signal, whereas the latter enables inclusion of more distal candidate genes. Any gene that meets at least one of these criteria is included in the list of bottom-up candidate genes. In addition, those candidate causal genes that contain either a sentinel or proxy variant of high or moderate impact on gene function (**Material and Methods**) are annotated as such.

#### Top-down component

For the top-down component, we first identify all genes that reside within a pre-defined genomic window either side of the sentinel variant. Various open-source databases are then referenced to determine whether any of these genes have previously been implicated in the regulation of the molecular phenotype in question, thereby constituting the top-down candidates (**Figure 1**). The type of databases referenced, and the way in which they are queried (**Supplementary Table S3**), depends on the nature of the molecular phenotype (e.g., the abundance of proteins, metabolites, lipids, etc.). For the purposes of this study, we extracted a list of metabolic-related genes from five databases: (i) Gene Ontology (GO), (ii) Kyoto Encyclopedia of Genes and Genomes (KEGG), (iii) Mouse Genome Informatics (MGI), (iv) Orphanet, and (v) Reactome (**Figure 1**; **Supplementary Table S3**). We have made this list available at GitHub (https://github.com/ds763/ProGeM). Lastly, the top-down candidate genes are assigned a score ranging between 1 and 5 to reflect the number of times they are reported in the databases.

#### Framework integration

The lists of bottom-up and top-down candidate genes for each identified QTL are integrated by ProGeM to determine whether any genes are identified by both independent approaches. Any concurrent candidate genes are then designated the most likely causal genes given that they fulfil both of the aforementioned requirements for a candidate causal gene.

### Generation of a high-confidence metabolite QTL reference causal gene set

In order to empirically assess the performance of ProGeM, we curated a reference dataset comprising 227 literature-derived metabolite QTLs (mQTLs), each of which we assigned a high-confidence causal gene. A full description on how this reference gene set was generated can be found in the **Material and Methods** section.

In brief, this reference set exploits the wealth of biochemical experimental research that predates GWAS discoveries, such as the identification and characterisation of proteins that regulate specific metabolic processes, as well as the extensive clinical characterisation of genes underlying rare inborn errors of metabolism. The candidate causal genes we assigned to these mQTLs affect the corresponding metabolites in a variety of ways; for example, many encode enzymes that act directly on the metabolite, others encode transporters or carriers for the metabolite, whilst others code for transcription factors known to impact the transcription of metabolic genes or processes. Full details including relevant enzyme commission (EC) codes and references (PubMed IDs) can be found in **Supplementary Table S1.** A summary and selection of representative examples are shown in **Table 1**.

**Table 1.** Summary of the biological relationships between expert-curated causal genes at 227 metabolite QTLs (mQTLs) and the corresponding metabolites. A selection of example mQTLs is included for illustrative purposes. The inclusion of “mQTL number” facilitates cross-referencing of example mQTLs between this table and Supplementary Table S1, where more detailed information can be found.

| Functional relationship (curated causal gene: metabolite) |  | Number of curated causal genes |  | Metabolite QTL (mQTL) example |  |  |  |
| --- | --- | --- | --- | --- | --- | --- | --- |
|  |  |  |  | Sentinel variant | Metabolite | Curated causal gene |  |
|  |  | Count | % |  |  | HGNC symbol | Gene name |
| Enzyme impacting metabolite | Directly | 66 | 29.07 | rs532545 | Uridine | CDA | Cytidine deaminase |
|  |  |  |  | rs4738684 | Glycine | GLDC | Glycine dehydrogenase |
|  | Indirectly | 46 | 20.26 | rs1801133 | Homocysteine | MTHFR | Methylenetetrahydrofolate reductase |
|  |  |  |  | rs12785878 | Vitamin D | DHCR7 | 7-Dehydrocholesterol reductase |
| Transporter or carrier for metabolite |  | 38 | 16.74 | rs1776029 | Manganese | SLC30A10 | Solute carrier family 30, member 10 |
|  |  |  |  | rs10455872 | Total cholesterol | LPA | Lipoprotein, Lp(a) |
| Receptor or binding partner for metabolite |  | 10 | 4.41 | rs2366858 | HDL cholesterol | CD36 | CD36 molecule (thrombospondin receptor) |
|  |  |  |  | rs12150660 | Testosterone | SHBG | Sex hormone-binding globulin |
| Affects transcription of related metabolic gene |  | 13 | 5.73 | rs6048216 | Fasting glucose | FOXA2 | Forkhead box A2 |
|  |  |  |  | rs603424 | Thyroid hormone levels (FT4) | LHX3 | LIM homeobox 3 |
| Acts on molecule / pathway known to impact metabolite |  | 21 | 9.25 | rs2954022 | Triglycerides | TRIB1 | Tribbles pseudokinase 1 |
|  |  |  |  | rs1801725 | Calcium | CASR | Calcium-sensing receptor |
| Acts on related metabolite |  | 29 | 12.78 | rs646776 | LDL cholesterol | SORT1 | Sortilin 1 |
|  |  |  |  | rs3738934 | X-13431-nonanoylcarnitine | ACADL | Acyl-CoA dehydrogenase, long chain |
| Other |  | 4 | 1.76 | rs5030062 | bradykinin, des-arg(9) | KNG1 | Kininogen 1 |
|  |  |  |  | rs38855 | Triglycerides | CAV1 | Caveolin 1 |
| Total |  | 227 | 100% |  |  |  |  |

### ProGeM implementation and parameter selection

ProGeM is implemented in the R statistical environment as a configurable .R script, which is freely available at GitHub along with a .readme file describing the necessary input and resultant output files (https://github.com/ds763/ProGeM).

The parameters used by ProGeM can be adjusted based on the type of molecular QTL data and the research question provided by the user. Specifically, these parameters include (i) the number of nearest genes to each sentinel variant that ProGeM should consider to be candidate causal genes (“number of nearest genes”; default=3); (ii) the size of the genomic window around each sentinel variant from which candidate genes are reported (“distance”; default=500kb); (iii) the threshold ProGeM should use to select proxies from a user-supplied file (“r^2^ threshold”; default≥0.8); and (iv) the threshold ProGeM should use to select *cis*-eQTL target genes as candidate causal genes (“*cis*-eQTL *p*-value threshold”=default: beta distribution-adjusted empirical *p*-values using a FDR threshold of 0.05, see http://www.gtexportal.org/home/documentationPage for details).

In order to determine how the ProGeM output is affected by changing various parameters, we applied ProGeM to the above-mentioned mQTL reference causal gene set (**Material and Methods; Supplementary Table S1**). We ran an additional iteration of ProGeM after each parameter change (while leaving all others in the default setting), and then determined both sensitivity and specificity. A high sensitivity is achieved if the identified sets of candidate causal genes include the “true positive” causal genes at the molecular QTLs, and a high specificity is obtained if the number of identified genes that do not match the “true positive” causal genes is low. Overall, we found that there was very little variation in either of these two metrics (**Supplementary Figure S1**), indicating that the general performance of ProGeM is robust to parameter changes.

Using the default parameters, we applied ProGeM to the set of 227 mQTLs for the purposes of performance benchmarking (see below). This analysis took approximately 15 minutes using a Windows 7 desktop equipped with an Intel^®^ core^TM^ i3-3240 (3.4GHz) processor and 4GB RAM. The bottom-up, top-down, and concurrent ProGeM outputs for this set of mQTLs can be found in **Supplementary Tables S4**, **S5**, and **S6**, respectively.

### Application and benchmarking of ProGeM

#### Local benchmarking

To illustrate specific characteristics of the ProGeM output in more detail, we arbitrarily selected three sentinel variants (rs1801133, rs1005390, and rs766420) (**Table 2**) from the full list of 227 mQTLs (**Supplementary Table S1**).

**Table 2.** Summary of the ProGeM output for three arbitrarily selected mQTLs. All bottom-up, top-down, and concurrent candidate causal genes are shown, as well as the implicating data sources. The expert-curated causal candidate genes are highlighted in red. <u>Abbreviations</u>: Chr; Chromosome, LD; Linkage Disequilibrium, GO; Gene Ontology, KEGG; Kyoto Encyclopedia of Genes and Genomes, MGI; Mouse Genome Informatics, Orph; Orphanet, Reac; Reactome.

| Annotation |  |  |  |  | Bottom-up |  |  | Top-down |  |  |  |  |  | Concurrent genes |
| --- | --- | --- | --- | --- | --- | --- | --- | --- | --- | --- | --- | --- | --- | --- |
| | Sentinel rsID | Chr | Position (GRCh37) | Number proxies ($r^2 \geq 0.8$) | LD region overlapping genes | Nearest gene(s) | Cis-eQTL target gene(s) | GO | KEGG | MGI | Orph | Reac | Top-scoring top-down candidate(s) | |
| 1 | rs1801133 | 1 | 11856378 | 0 | *MTHFR* | *MTHFR*; *C1orf167*; *CLCN6* | *MFN2*; *MTHFR* | *MTHFR*; *NPPA*; *PTCHD2* | *MTHFR* | *MAD2L2*; *NPPA* | *MTHFR* | *MTHFR* | *MTHFR* (4) | *MTHFR* |
| 107 | rs1005390 | 7 | 150543721 | 26 | *AOC1* | *AOC1*; *TMEM179A*; *TMEM179B* | *AOC1*; *TMEM179A*; *TMEM179B* | *AOC1*; *SMARCD3* | *AOC1*; *CHPF2*; *NOS3* | *NOS3* | - | *AOC1*; *CHPF2*; *NOS3*; *SMARCD3* | *AOC1* (3); *NOS3* (3) | *AOC1*; *CHPF2* |
| 213 | rs766420 | X | 153554404 | 0 | *TKTL1* | *TKTL1*; *FLNA*; *TEX28* | *BRCC3*; *DNASE1L1*; *F8A1*; *FAM3A*; *FAM50A*; *GDI1*; *IRAK1*; *LAGE3*; *PLXNA3*; *RPL10*; *TAZ*; *TMEM187* | *DNASE1L1*; *G6PD*; *IDH3G*; *MECP2*; *MPP1*; *RENBP*; *TKTL1* | *ATP6AP1*; *G6PD*; *IDH3G*; *TKTL1* | *DKC1*; *IKBKG* | *HCFC1*; *SSR4*; *TAZ* | *G6PD*; *IDH3G*; *RPL10*; *TAZ* | *G6PD* (3); *IDH3G* (3) | *DNASE1L1*; *RPL10*; *TKTL1* |

#### Example 1: rs1801133 and homocysteine

rs1801133 has been previously identified to associate with plasma homocysteine levels by two large-scale GWAS at genome-wide significance (34,35). rs1801133 is a missense variant (Ala222Val) affecting the *MTHFR* gene, which encodes an enzyme known to be involved in folate and homocysteine metabolism (36). Therefore, *MTHFR* was assigned to this mQTL as the high-confidence causal gene (**Table 2**). Accordingly, the ProGeM output identified *MTHFR* as the sole concurrent candidate causal gene at this mQTL, highlighting the expert-curated gene as the most likely causal gene. Indeed, the bottom-up component of ProGeM indicated that *MTHFR* is the nearest protein-coding gene to rs1801133 as well as the only gene overlapping the LD region, though it was not highlighted as a *cis*-eQTL gene. Further, *MTHFR* was implicated by four out of the five top-down databases probed – the Mouse Genome Informatics (MGI) database being the exception.

#### Example 2: rs1005390 and X-03056—N-[3-(2-Oxopyrrolidin-1-yl)propyl]acetamide

In a GWAS investigating 529 blood metabolites, rs1005390 was found to be significantly (*p*=2.61×10^−16^) associated with circulating X-03056—N-[3-(2-Oxopyrrolidin-1-yl)propyl]acetamide levels (17). This variant is intronic to the high-confidence causal gene, *AOC1* (**Table 2**), which encodes an enzyme that catalyses the deamination of N1-acetylspermidine to produce the above-mentioned metabolite (37). As was the case with example 1, the expert-curated gene was identified by ProGeM as a concurrent candidate causal gene, although in this case, *AOC1* was accompanied by a second concurrent gene, *CHPF2*. Of the two genes, *AOC1* was highlighted by all three of the bottom-up data sources as well as three top-down databases. Furthermore, *AOC1* was annotated as the nearest protein-coding gene to rs1005390 whereas *CHPF2* resides almost 400kb downstream, which suggests that the nearest gene information may be useful when prioritising between concurrent candidates.

#### Example 3: rs766420 and bilirubin

In a GWAS investigating the genetic determinants of circulating levels of bilirubin, which is a by-product of the breakdown of haemoglobin in red blood cells, the authors reported a significant (*p*=9.40×10^−9^) association with rs766420 (38). This variant resides within an intron of the gene *TKTL1*. In contrast to the other two examples above, the high-confidence causal gene, in this case, was not the most proximal gene but rather the gene *G6PD* (**Table 2**), which is located more than 200kb downstream. *G6PD* encodes glucose-6-phosphate dehydrogenase, an enzyme that is critical for red blood cell metabolism, as deficiency is known to result in haemolysis, anaemia, hyperbilirubinemia, and jaundice (39). Although the ProGeM output for this mQTL highlighted three genes as concurrent candidates (i.e., *DNASE1L1*, *RPL10*, and *TKTL1*), none of them corresponded to the expert-curated gene, *G6PD*. *G6PD* was not highlighted by the bottom-up component of ProGeM; however, it did feature as a top-down candidate with supporting evidence from three databases. Taken together, this example underscores the importance of incorporating top-down information within the ProGeM framework, whilst also cautioning against automatically discounting non-concurrent genes without due diligence.

#### Global benchmarking

Following the characterisation of the ProGeM output at individual molecular QTLs, we next determined key performance indicators. In order to benchmark the sensitivity and specificity of ProGeM, we systematically compared the ProGeM output (stratified for the bottom-up and top-down component) for all 227 mQTLs with the corresponding high-confidence causal gene assignments as described above (**Material and Methods**).

To assess the sensitivity, we determined the proportion of reference candidate causal genes at the mQTLs (“true positives”) that were identified by ProGeM. Overall, ProGeM was able to identify the curated candidates for 223 of 227 mQTLs, thereby demonstrating high sensitivity (98%) (**Figure 2A**). Importantly, the vast majority of these genes (*n*=187; 82%) were identified by both the bottom-up and top-down components (**Figure 2B**), indicating that sensitivity remains high even when restricting to the narrower set of concurrent candidates. Indeed, the bottom-up (**Supplementary Figure S2**) and top-down (**Supplementary Figure S3**) components alone identified 216 (95%) and 194 (85%) true candidate causal genes, respectively.

**Figure 2.**
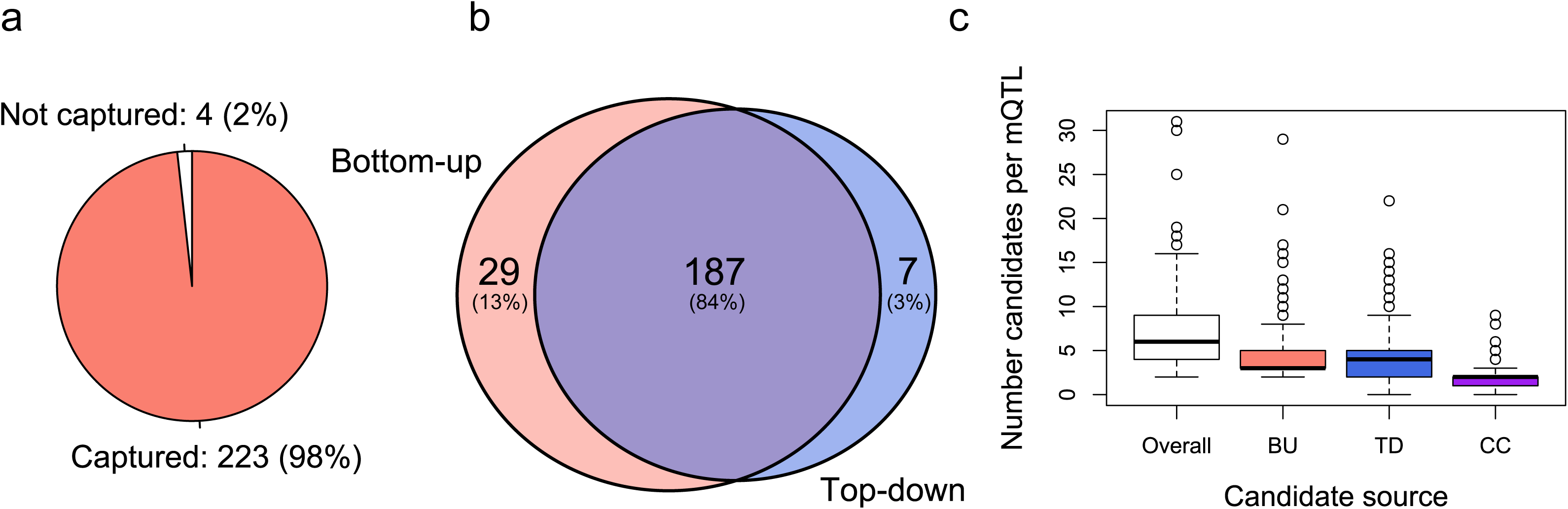
Global benchmarking of ProGeM using high-confidence causal gene assignments at 227 metabolite QTLs. **(a)** Number (percentage) of high-confidence candidate causal genes captured by our framework; **(b)** Summary of the number of high-confidence causal genes captured either uniquely or by both the bottom-up and top-down components of ProGeM; **(c)** Box plot summarising the number of candidate causal genes identified for each metabolite QTL by our framework overall (total), as well the bottom-up (BU) and top-down (TD) components uniquely and concurrently (CC). The box plot shows the median and interquartile ranges, with the whiskers extending to 1.5-times the corresponding interquartile range. Data points outside of this range are indicated individually as circles.

Next, we assessed the specificity of ProGeM across all 227 mQTLs. In total, ProGeM highlighted 1,629 candidate causal genes at these loci, with a median of 6 [min=2, max=31] candidates per locus (**Figure 2C**). The overall specificity of ProGeM for this dataset was 0.502 (**Material and Methods**). When we compared the bottom-up and top-down components of ProGeM, we found that the top-down component performed slightly better with a specificity of 0.459 compared to 0.384 for the bottom-up component. Nevertheless, the two components had very similar medians and ranges (**Figure 2C**; **Supplementary Figure S4**). Notably, when only the concurrent candidate causal genes were taken into account, specificity was much improved with an overall specificity of 0.846, corresponding to a median of just 2 [min=0, max=9] candidate at each locus (**Figure 2C**).

### Comparative analysis of functional annotation data sources

One of the main limitations of current bioinformatic tools for prioritising candidate causal genes at molecular QTL, including ProGeM, pertains to the difficulties associated with distinguishing between true and false positive candidates. Our benchmarking analyses showed that focusing solely on the concurrent gene set returned by ProGeM appears to be a potential means to address this issue. To formally test this, we performed an enrichment analysis using all non-concurrent candidates highlighted by ProGeM for the same mQTL data (*n*=227 loci) as a background set. A Fisher’s exact test revealed a highly significant odds ratio (OR) of 28.55 [95% confidence interval: 19.26-43.19] (*p*=8.47×10^−93^), indicating that the odds of identifying the true causal genes from candidates highlighted by ProGeM are vastly improved when picking from the pool of concurrent candidate genes. Indeed, based on the observed frequencies, 46% of the concurrent candidates corresponded to a reference causal gene, relative to only 3% of non-concurrent candidate genes (**Supplementary Table S7**).

Next, we assessed the importance of the various functional data sources leveraged by ProGeM for pinpointing the true positive candidate causal genes. Of all the bottom-up annotation criteria tested, the set of genes nearest to a sentinel variant was enriched with true positive candidate causal genes at the highest significance level (OR=45.09 [30.66-67.23], *p*=1.02×10^−107^) (**Figure 3**; **Supplementary Table S7**). This was followed by the concurrent gene set (see above), the LD overlapping gene set (OR=15.17 [10.61-22.02], *p*=9.48×10^−67^), and then the three nearest genes to a sentinel variant (OR=14.51 [9.44-23.10], *p*=8.19×10^−55^) (**Figure 3**; **Supplementary Table S7**). Given that three of the four most significant criteria relate to the most proximal genes to the sentinel variant at the mQTLs, it can be concluded that proximity-based criteria are effective indicators of true positive causal genes at mQTLs.

**Figure 3.**
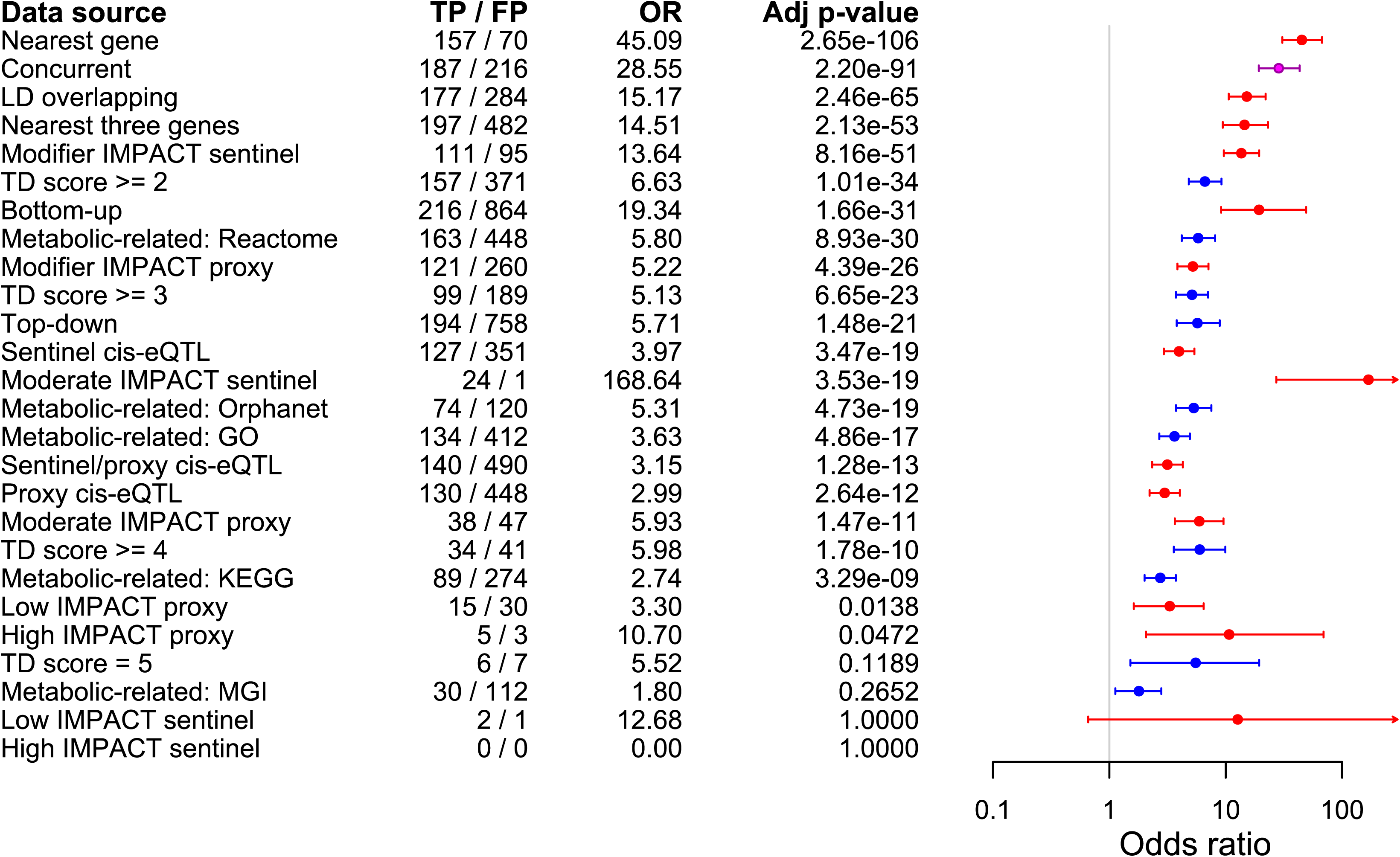
Comparison of top-down and bottom-up functional criteria for distinguishing true from false positive mQTL causal gene candidates. Odds ratios and 95% confidence intervals indicating the likelihood that candidate causal genes with various bottom-up or top-down characteristics correspond to the high-confidence causal gene assignments (i.e., true causal genes). The background gene set for enrichment analysis of each characteristic was comprised of all remaining candidates identified by ProGeM. Fisher’s exact test was used throughout, and Bonferroni-corrected (26 tests) *p*-values are indicated. The number of true positive (TP) and false positive (FP) causal genes identified by each characteristic are also indicated. The boxes and confidence intervals are colour-coded according to whether they correspond to bottom-up (red), top-down (blue), or concurrent (purple) data sources.

When we compared the concurrent and nearest gene sets more closely, the concurrent gene set achieved higher sensitivity, having identified 187 (82%) relative to 157 (69%) true positive causal genes out of 227. However, the concurrent gene set exhibited lower specificity (0.846) than the nearest gene set, which inherently achieved maximal specificity. Given this, we constructed a combined gene set by imposing maximal specificity onto our concurrent gene set using nearest gene information as follows: (i) for mQTLs assigned to more than one concurrent candidate gene, we restricted this assignment to the concurrent gene nearest to the sentinel variant, and (ii) we assigned mQTLs without a concurrent candidate to the gene nearest to the sentinel variant. The resultant set of “nearest-concurrent” genes identified 177 of 227 (78%) reference causal genes, which constitutes a 4% drop in sensitivity relative to our original set of concurrent genes (187; 82%) but a 9% increase relative to the nearest gene set (157; 69%). Further, an enrichment analysis of the nearest-concurrent gene set revealed both a higher odds ratio and *p*-value (OR=84.10 [54.98-130.66], *p*=1.56×10^−137^) relative to both the original gene sets.

The annotation criterion with by far the highest odds ratio (OR=168.64 [27.20-6666.66], *p*=1.36×10^−20^) was related to genes containing a sentinel variant of moderate impact (**Figure 3**; **Supplementary Table S7**). Indeed, of 25 such candidate causal genes highlighted by ProGeM, 24 (96.0%) corresponded to a reference causal candidate (**Supplementary Table S7**). However, although genes containing a proxy (r^2^>0.8) variant of moderate impact were also enriched with true positive causal genes, this gene set was associated with a much lower odds ratio (OR=5.93 [3.66-9.57], *p*=5.64×10^−13^) (**Figure 3**; **Supplementary Table S7**).

Because genes containing a sentinel variant of moderate impact will also inherently be the nearest gene to that sentinel, we sought to determine whether the highly significant enrichment observed for the nearest gene set was driven by the genes that contain a sentinel variant of moderate impact. Therefore, we repeated the enrichment analysis of the nearest gene set after removing all mQTLs tagged by a moderate impact sentinel variant, which resulted in a dataset that comprised 202 out of 227 mQTLs. There were no mQTLs tagged by a high impact sentinel in this dataset. We found that this filtered nearest gene set was still significantly enriched with true positive causal genes (OR=34.24 [23.03-51.38], *p*=1.33×10^−84^) (**Supplementary Table S8**), indicating that the enrichment observed for the complete nearest gene set (OR=45.09 [30.66-67.23], *p*=1.02×10^−107^) was not wholly driven by the genes containing moderate impact sentinel variants. Accordingly, we obtained comparable results when we also removed mQTLs tagged by a low impact sentinel variant as well as after removing mQTLs tagged by proxy variants of high, moderate, and low impact (**Supplementary Table S8**).

The *cis*-eQTL gene sets were also significantly enriched with true positive causal genes, although the *p*-values and associated odds ratios were modest (**Figure 3**; **Supplementary Table S7**). When we investigated in more detail the 24 mQTLs for which the true positive causal gene contained a moderate impact sentinel variant, we found that just nine of these genes were also *cis*-eQTL genes for either the sentinel or a proxy variant. This indicates that these two methods of identifying true positive causal genes are predominantly exclusive. We also performed enrichment analyses for all GTEx tissues (*n*=48) assayed individually; however, we did not identify any specific tissues of particular relevance (**Supplementary Figure S5; Supplementary Table S9**). The same applied to individual top-down annotation criteria tested (**Figure 3**; **Supplementary Table S7**).

### Cross-validation and partial replication in a large-scale pQTL dataset

Next, we assessed the extent to which these functional annotation criteria might serve as indicators of true positive causal genes in another molecular QTL dataset. To this end, we utilised a *cis*-pQTL dataset comprising 562 sentinel variants (14), for which we hypothesised that the true positive causal gene ought to be the gene that encodes the *cis*-affected protein (**Material and Methods**; **Supplementary Table S2**). Importantly, this dataset only enabled us to assess the bottom-up annotation criteria, as the top-down criteria are not directly comparable across the mQTL and pQTL datasets. Therefore, we used only the bottom-up candidate genes from the corresponding ProGeM output (obtained using default settings) as our background gene set. Likewise, for the purposes of this comparison, we also re-ran the enrichment analyses of the bottom-up criteria for the mQTL dataset, where we used only the bottom-up candidates highlighted by ProGeM as a background gene set.

The findings for both the mQTL (**Supplementary Table S10**) and *cis*-pQTL (**Supplementary Table S12**) datasets (**Figure 4**) were strikingly similar. First, not only did the set of nearest genes achieve the highest levels of significance in both cases (*cis*-pQTL: OR=53.59 [40.51-71.19], *p*=4.13×10^−251^ | mQTL: OR=30.00 [20.14-45.33], *p*=2.40×10^−82^), but the other two proximity-based criteria (LD overlapping, nearest three genes) also appeared in the top four of their respective lists (ranked by *p*-value) (**Figure 4**). This is consistent with previous observations made for this *cis*-pQTL dataset, whereby the sentinel variants were found to cluster at the transcription start site (TSS) of genes encoding the *cis*-proteins (14). Second, the gene sets containing a sentinel variant of moderate impact achieved the highest odds ratios for both datasets (*cis*-pQTL: OR=99.37 [37.22-373.58], *p*=4.64×10^−58^ | mQTL: OR=107.35 [17.29-4326.68], *p*=1.19×10^−16^) (**Figure 4**). Third, the significant enrichment observed within the nearest gene sets from either dataset was only partially attenuated after removing QTLs tagged by these moderate impact sentinel variants (**Supplementary Tables S8**; **S12**). These data indicate that genomic proximity to the sentinel variant represents a strong indicator of the true positive causal genes for both *cis*-pQTLs and mQTLs – even if the variant in question does not reside in the coding sequence.

**Figure 4.**
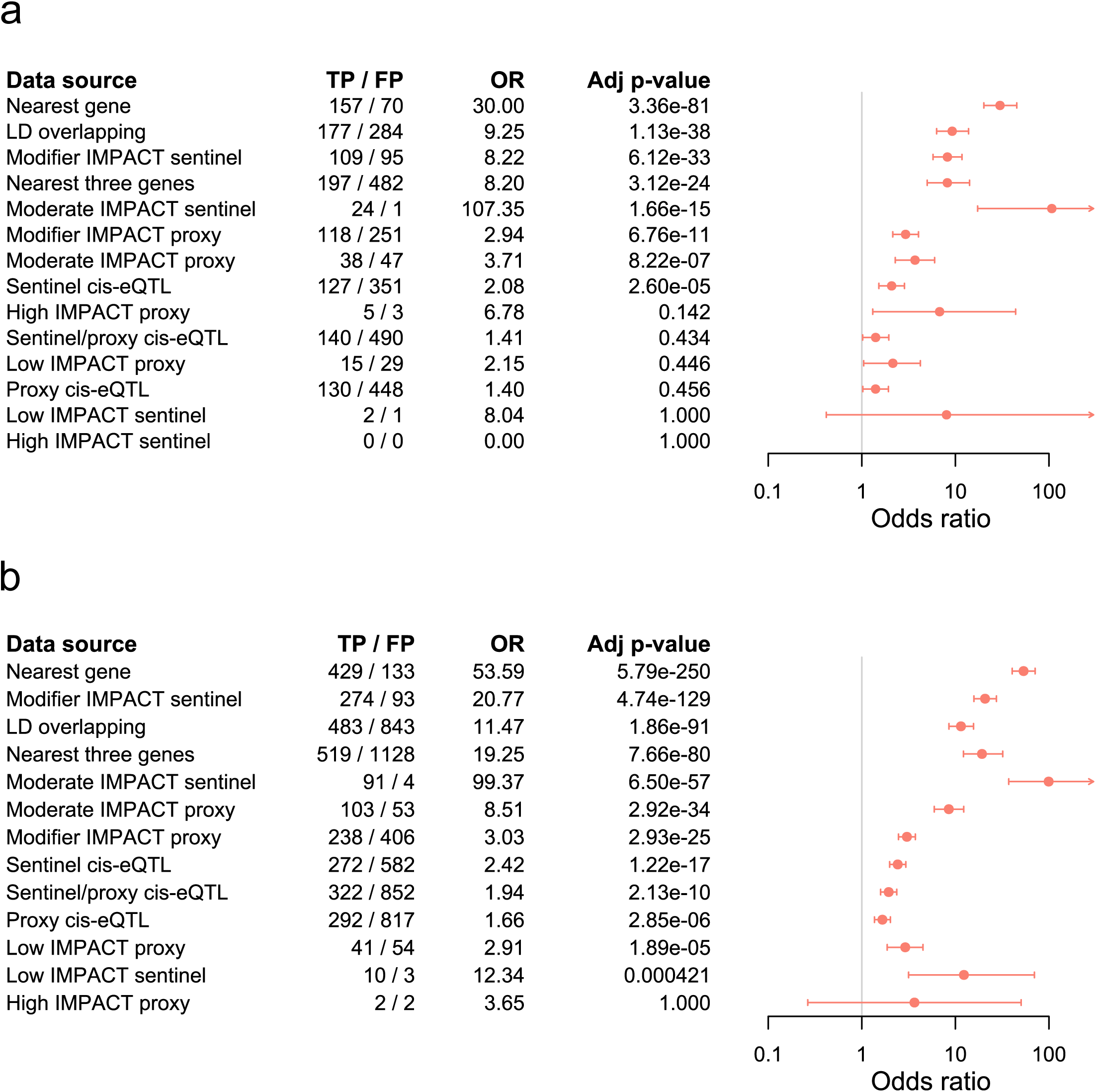
Comparison of bottom-up functional criteria for distinguishing true from false positive (a) *cis*-pQTL and (b) mQTL causal gene candidates. Odds ratios and 95% confidence intervals indicating the likelihood that candidate causal genes with various bottom-up characteristics correspond to the high-confidence causal genes (i.e., true causal genes). The background gene set for enrichment analysis of each characteristic was comprised of all remaining bottom-up candidates identified by ProGeM for the relevant ‘omic modality. Fisher’s exact test was used throughout, and Bonferroni-corrected (14 tests) *p*-values are indicated. The number of true positive (TP) and false positive (FP) causal genes identified by each characteristic are also indicated.

Enrichment analyses of the *cis*-eQTL target gene sets yielded similar odds ratios across the two datasets, although a higher significance was observed for the *cis*-pQTL dataset (**Supplementary Tables S10**; **S11**). Further, when we performed enrichment analyses of each GTEx tissue individually, both the odds ratios and *p*-values observed for the *cis*-pQTL dataset were generally more significant than for the mQTL dataset (**Supplementary Figures S6**; **S7**; **Supplementary Tables S9**; **S13**). However, this difference in *p*-values may be due to the fact that the *cis*-pQTL dataset comprised more than twice the number of QTLs relative to the mQTL dataset (i.e., 562 vs. 227 QTLs), thereby leveraging greater statistical power.

Taken together, these results suggest that many of the same bottom-up criteria can be used to effectively prioritise the most likely true positive causal genes for both mQTLs and pQTLs, and that ProGeM may be applicable to other molecular QTL datasets beyond those tested here.

## DISCUSSION

In the present study, we introduced an analysis framework and highly configurable R script for the **Pr**ioritisation **o**f candidate causal **Ge**nes at **M**olecular QTLs (ProGeM). In benchmarking analyses using a set of 227 mQTLs for which we mapped high-confidence causal genes, we demonstrated that ProGeM is highly sensitive in identifying these genes. Further, in enrichment analyses using this mQTL dataset, we found that proximity-based indicators are an effective means of distinguishing between true and false positive causal genes.

### Unique features of ProGeM

First, currently available tools are typically either trait agnostic or aimed more generally at complex disease GWAS data, whereas ProGeM is intended for a specific trait class, i.e., molecular QTLs. This confers a major advantage, as knowledge of the trait inherently enables the incorporation of trait-specific annotation criteria. In the present study, we demonstrated this for “metabolism” as a broad trait class, whereby we extracted metabolic-related genes from five freely available databases (**Material and Methods**) and cross-referenced them against genes in proximity to each corresponding mQTL sentinel variant. We note that there are other web tools available that utilise top-down information; however, these tools are distinct in that they rely on either user-input (i.e., Phenolyzer (40)) or literature/text mining (i.e., PolySearch (41), MimMiner (42), Bitola (43), aGeneApart (44), GeneProspector (45)) to generate this information.

Second, ProGeM integrates both bottom-up and top-down functional annotation data sources to hone in specifically on “concurrent” candidate causal genes. In our benchmarking tests, we demonstrated that the concurrent candidate causal genes for a set of 227 mQTLs were strongly enriched with the high-confidence causal genes. These data suggest that there is a benefit in prioritising concurrent over non-concurrent candidates. Furthermore, this benchmarking also revealed multiple cases for which the expert-curated candidate was highlighted by either bottom-up or top-down evidence, which indicates that the incorporation of two complementary layers of information is an effective means of maximising the sensitivity of ProGeM. Indeed, we observed this scenario in the example of the rs766420-bilirubin QTL discussed above, for which the reference causal gene, *G6PD*, which resides ~200kb from the corresponding sentinel variant, was prioritised by the top-down component of ProGeM but not the bottom-up component.

Third, ProGeM has been benchmarked against a large reference set of mQTLs with high-confidence causal gene assignments, which allowed us to empirically verify the validity and performance of our framework. By contrast, the published tools that have been benchmarked (e.g., ToppGene Suite) have used very small datasets of known causal genes, which are most likely not generalisable (46,47). Further, benchmarking of the majority of available tools has not incorporated a baseline to empirically assess performance, but rather has comprised the application of the tool to a complex disease GWAS dataset followed by a description of the output (e.g., FUMA (20)). This paucity of systematic benchmarking against a high-confidence reference for candidate causal gene prioritisation tools is due to the fact that appropriate and large reference gene sets have been difficult to generate for molecular QTLs. We have made our mQTL reference dataset available to the research community in **Supplementary Table S1**, providing a substrate for future benchmarking analyses and methods development.

### Global benchmarking analyses

Our benchmarking analyses highlighted increased sensitivity to be one of the strengths of ProGeM, having missed only four out of 227 reference causal genes at the tested mQTLs (**Figure 2**). When we investigated these four elusive genes in more detail, we found that three of them were missed because they are located >500kb from their respective sentinel variants (rs7542172; *AKR1A1* | rs140348140; *GLDC* | rs1550532; *TRPM8*), whilst the fourth was missed because it was annotated as a pseudogene (rs7130284; *FOLH1B*). This potentially explains why ProGeM was unable to capture the curated causal gene for rs10403668. Nevertheless, we were able to achieve maximal sensitivity by modifying two user-defined settings in ProGeM as follows: (i) omit the default filter on protein-coding genes only, and (ii) extend the genomic locus from the default +/-500kb to +/-1mb.

It is important to note that the high sensitivity achieved by ProGeM was inevitably accompanied by high levels of “background noise” (i.e., low specificity; **Figure 2**). This is not unusual within the context of candidate causal gene prioritisation tools, whereby the general focus tends to be on “prioritising” multiple candidates at a locus rather than force-assigning each QTL to a single causal candidate – notwithstanding that at some molecular QTLs there may be more than one causal gene. Thus, in order to minimise the background noise associated with candidate causal gene prioritisation tools, there is a general need to be able to apply additional prioritisation strategies post-hoc that are both reliable and empirically validated. For example, as mentioned above, our comprehensive benchmarking analyses demonstrated that by specifically prioritising the concurrent candidate causal genes highlighted by ProGeM, we were able to make considerable specificity gains at the cost of only a minimal reduction in sensitivity.

Our analyses also showed that genomic proximity to the sentinel variants tagging mQTLs was a highly effective means of prioritising candidates. Indeed, out of multiple candidate gene sets defined by a series of bottom-up and top-down criteria, the set of genes nearest to each sentinel variant achieved the highest level of significance in enrichment testing, with the concurrent gene set ranking second (**Figure 3**). Furthermore, the subclass of nearest genes that contained a moderate impact sentinel variant achieved the highest odds ratio of all criteria tested, whereby out of 25 such genes in total, 24 corresponded to a true positive causal gene. Although this relates to a relatively small number of genes from the full dataset of 227 causal genes, it suggests that genes containing moderate impact sentinel variants are reliable indicators of true causal genes at mQTLs. Importantly, we also showed that although the enrichment signal observed for the full set of nearest genes was enhanced by the subset of genes harbouring moderate impact sentinel variants, they did not drive this signal. Thus, even in the absence of these genes, the nearest gene set still served as a reliable indicator of true positive causal genes at mQTLs.

In contrast, the general consensus in recent years has been that the underlying genetic risk factors for complex human diseases and traits are primarily regulatory in nature, whilst the nearest gene to a sentinel variant often does not correspond to the true causal gene (48). Notably, both the mQTL and *cis*-pQTL datasets highlighted *cis*-eQTL target genes (as reported in GTEx v7 data) as relatively poor indicators of “true positive” causal genes. Further research is needed to assess whether genomic proximity is a good indicator of a true positive causal gene for other molecular QTLs as well as complex disease traits, and will depend on the availability of high-confidence reference datasets.

### Current limitations of ProGeM

A potential limitation of ProGeM relates to the preparation of the reference causal gene assignments at the 227 metabolite QTLs, which could have been subject to bias, such as the prioritisation of the nearest genes for the assignments. As outlined in detail in the **Material and Methods** section, all genes within 1mb of the sentinel variants were included for the annotation and in-depth literature review. In support, our findings obtained using the bottom-up component of ProGeM were consistent across both the mQTL and *cis*-pQTL datasets. The latter reference set was derived using distinct methods (**Materials and Methods**). This suggests that our conclusion, that genomic proximity to the sentinel variant is a reliable indicator of a true positive causal candidate gene at molecular QTLs, has validity.

Furthermore, although the overall approach employed by ProGeM and the methods used to curate the high-confidence mQTL dataset were different (**Material and Methods**), it is possible that some of the databases utilised by ProGeM may have been informed by the biochemical literature used for the expert curation. Therefore, we acknowledge that ProGeM and the curated high-confidence mQTL dataset are not entirely independent of each other, and as a result, the sensitivity of ProGeM observed within the context of mQTL may be inflated. We also note that the KEGG database was utilised both by ProGeM and as a guide for the expert literature review, although KEGG was just one of five top-down databases utilised by ProGeM. Our data showed that this did not result in a biased annotation.

### Possible extensions of ProGeM

Looking ahead, we anticipate the application of ProGeM to molecular QTL datasets from additional ‘omic modalities in the future. This may be directly applicable for lipid QTLs due to the similarities of standardised assay platforms but less so for *trans*-pQTLs, for example. Indeed, the prioritisation of candidate causal genes at *trans*-pQTLs would call for a different top-down strategy to the one adopted here. Accordingly, we have previously applied a “guilt-by-association” (GbA) strategy towards the annotation of *trans*-pQTLs (14). Thus, rather than asking whether genes local to a sentinel variant have previously been implicated in a metabolic-related phenotype, we asked whether any local genes exhibit related functioning to the gene encoding a given *trans*-affected protein; i.e., annotation within the same biological pathway, or evidence of a protein-protein interaction (PPI). Notably, multiple currently available tools intended for complex human disease have also adopted GbA strategies. These approaches work under the assumption that unknown or novel causal genes can be identified on the basis that they must exhibit related functionality to known causal genes (47,49). There is, therefore, ample precedence for GbA within the context of candidate causal gene prioritisation.

### Conclusions

In summary, ProGeM is a new gene prioritisation tool developed specifically for the identification and prioritisation of candidate causal genes at molecular QTLs. We have demonstrated its utility for mQTLs, with one of its major strengths being high sensitivity. We have also highlighted multiple criteria that can be used to prioritise certain candidates over others at a given mQTL. Within the ProGeM framework, we provided evidence that those candidate causal genes with both bottom-up and top-down supporting evidence (i.e., concurrent candidates) represent likely true causal genes. We also showed that proximity to the sentinel variant is a reliable indicator of a true positive causal gene, particularly those genes containing a sentinel variant of moderate impact (i.e., missense variants). Based on our findings, we caution against an overreliance on *cis*-eQTL target genes, as it appears that long-range regulatory effects at molecular QTLs appear may be the exception rather than the rule.

## AVAILABILITY

ProGeM is freely available in the form of an R script at the GitHub repository (https://github.com/ds763/ProGeM).

## ACKNOWLEDGEMENT

We thank Parsa Akbari, William Astle, Jessica Barrett, James Blackshaw, Stephen Burgess, Qi Guo, Joanna Howson, Tao Jiang, Mihir Kamat, Clare Oliver-Williams, James Peters, Bram Prins, Jessica Rees, and Praveen Surendran for helpful comments on the analysis framework.

## REFERENCES

1. MacArthur, J., Bowler, E., Cerezo, M., Gil, L., Hall, P., Hastings, E., Junkins, H., McMahon, A., Milano, A., Morales, J. et al. (2017) The new NHGRI-EBI Catalog of published genome-wide association studies (GWAS Catalog). Nucleic Acids Res, 45, D896–D901.

2. Nelson, C.P., Goel, A., Butterworth, A.S., Kanoni, S., Webb, T.R., Marouli, E., Zeng, L., Ntalla, I., Lai, F.Y., Hopewell, J.C. et al. (2017) Association analyses based on false discovery rate implicate new loci for coronary artery disease. Nat Genet, 49, 1385–1391.

3. Ripke, S., Neale, B.M., Corvin, A., Walters, J.T., Farh, K.H., Holmans, P.A., Lee, P., Bulik-Sullivan, B., Collier, D.A., Huang, H. et al. (2014) Biological insights from 108 schizophrenia-associated genetic loci. Nature, 511, 421–427.

4. Visscher, P.M., Wray, N.R., Zhang, Q., Sklar, P., McCarthy, M.I., Brown, M.A. and Yang, J. (2017) 10 Years of GWAS Discovery: Biology, Function, and Translation. Am J Hum Genet, 101, 5–22.

5. Ardlie, K.G., Deluca, D.S., Segre, A.V., Sullivan, T.J., Young, T.R., Gelfand, E.T., Trowbridge, C.A., Maller, J.B., Tukiainen, T., Lek, M. et al. (2015) Human genomics. The Genotype-Tissue Expression (GTEx) pilot analysis: multitissue gene regulation in humans. Science, 348, 648–660.

6. Cookson, W., Liang, L., Abecasis, G., Moffatt, M. and Lathrop, M. (2009) Mapping complex disease traits with global gene expression. Nat Rev Genet, 10, 184–194.

7. Dixon, A.L., Liang, L., Moffatt, M.F., Chen, W., Heath, S., Wong, K.C., Taylor, J., Burnett, E., Gut, I., Farrall, M. et al. (2007) A genome-wide association study of global gene expression. Nat Genet, 39, 1202–1207.

8. Montgomery, S.B., Sammeth, M., Gutierrez-Arcelus, M., Lach, R.P., Ingle, C., Nisbett, J., Guigo, R. and Dermitzakis, E.T. (2010) Transcriptome genetics using second generation sequencing in a Caucasian population. Nature, 464, 773–777.

9. Stranger, B.E., Nica, A.C., Forrest, M.S., Dimas, A., Bird, C.P., Beazley, C., Ingle, C.E., Dunning, M., Flicek, P., Koller, D. et al. (2007) Population genomics of human gene expression. Nat Genet, 39, 1217–1224.

10. Bonder, M.J., Luijk, R., Zhernakova, D.V., Moed, M., Deelen, P., Vermaat, M., van Iterson, M., van Dijk, F., van Galen, M., Bot, J. et al. (2017) Disease variants alter transcription factor levels and methylation of their binding sites. Nat Genet, 49, 131–138.

11. Wahl, S., Drong, A., Lehne, B., Loh, M., Scott, W.R., Kunze, S., Tsai, P.C., Ried, J.S., Zhang, W., Yang, Y. et al. (2017) Epigenome-wide association study of body mass index, and the adverse outcomes of adiposity. Nature, 541, 81–86.

12. Lourdusamy, A., Newhouse, S., Lunnon, K., Proitsi, P., Powell, J., Hodges, A., Nelson, S.K., Stewart, A., Williams, S., Kloszewska, I. et al. (2012) Identification of cis-regulatory variation influencing protein abundance levels in human plasma. Hum Mol Genet, 21, 3719–3726.

13. Suhre, K., Arnold, M., Bhagwat, A.M., Cotton, R.J., Engelke, R., Raffler, J., Sarwath, H., Thareja, G., Wahl, A., DeLisle, R.K. et al. (2017) Connecting genetic risk to disease end points through the human blood plasma proteome. Nat Commun, 8, 14357.

14. Sun, B.B., Maranville, J.C., Peters, J.E., Stacey, D., Staley, J.R., Blackshaw, J., Burgess, S., Jiang, T., Paige, E., Surendran, P. et al. (2018) Genomic atlas of the human plasma proteome. Nature, 558, 73–79.

15. Stegemann, C., Pechlaner, R., Willeit, P., Langley, S.R., Mangino, M., Mayr, U., Menni, C., Moayyeri, A., Santer, P., Rungger, G. et al. (2014) Lipidomics profiling and risk of cardiovascular disease in the prospective population-based Bruneck study. Circulation, 129, 1821–1831.

16. Adamski, J. and Suhre, K. (2013) Metabolomics platforms for genome wide association studies-linking the genome to the metabolome. Curr Opin Biotechnol, 24, 39–47.

17. Shin, S.Y., Fauman, E.B., Petersen, A.K., Krumsiek, J., Santos, R., Huang, J., Arnold, M., Erte, I., Forgetta, V., Yang, T.P. et al. (2014) An atlas of genetic influences on human blood metabolites. Nat Genet, 46, 543–550.

18. Suhre, K., Shin, S.-Y., Petersen, A.-K., Mohney, R.P., Meredith, D., Wägele, B., Altmaier, E., CardioGram, Deloukas, P., Erdmann, J. et al. (2011) Human metabolic individuality in biomedical and pharmaceutical research. Nature, 477, 54.

19. Arnold, M., Raffler, J., Pfeufer, A., Suhre, K. and Kastenmuller, G. (2015) SNiPA: an interactive, genetic variant-centered annotation browser. Bioinformatics, 31, 1334–1336.

20. Watanabe, K., Taskesen, E., van Bochoven, A. and Posthuma, D. (2017) Functional mapping and annotation of genetic associations with FUMA. Nat Commun, 8, 1826.

21. Kimoto, M., Whitley, G.S., Tsuji, H. and Ogawa, T. (1995) Detection of NG, NG-dimethylarginine dimethylaminohydrolase in human tissues using a monoclonal antibody. J Biochem, 117, 237–238.

22. Nguyen, L.N., Ma, D., Shui, G., Wong, P., Cazenave-Gassiot, A., Zhang, X., Wenk, M.R., Goh, E.L. and Silver, D.L. (2014) Mfsd2a is a transporter for the essential omega-3 fatty acid docosahexaenoic acid. Nature, 509, 503–506.

23. Nielsen, J., Christiansen, J., Lykke-Andersen, J., Johnsen, A.H., Wewer, U.M. and Nielsen, F.C. (1999) A family of insulin-like growth factor II mRNA-binding proteins represses translation in late development. Mol Cell Biol, 19, 1262–1270.

24. Higgins, M.J., Lecamwasam, D.S. and Galton, D.J. (1975) A new type of familial hypercholesterolaemia. Lancet, 2, 737–740.

25. Michaely, P., Li, W.P., Anderson, R.G., Cohen, J.C. and Hobbs, H.H. (2004) The modular adaptor protein ARH is required for low density lipoprotein (LDL) binding and internalization but not for LDL receptor clustering in coated pits. J Biol Chem, 279, 34023–34031.

26. Weiss, M.J., Cole, D.E., Ray, K., Whyte, M.P., Lafferty, M.A., Mulivor, R.A. and Harris, H. (1988) A missense mutation in the human liver/bone/kidney alkaline phosphatase gene causing a lethal form of hypophosphatasia. Proc Natl Acad Sci U S A, 85, 7666–7669.

27. Srivenugopal, K.S. and Ali-Osman, F. (1997) Activity and distribution of the cysteine prodrug activating enzyme, 5-oxo-L-prolinase, in human normal and tumor tissues. Cancer Lett, 117, 105–111.

28. Leyva-Illades, D., Chen, P., Zogzas, C.E., Hutchens, S., Mercado, J.M., Swaim, C.D., Morrisett, R.A., Bowman, A.B., Aschner, M. and Mukhopadhyay, S. (2014) SLC30A10 is a cell surface-localized manganese efflux transporter, and parkinsonism-causing mutations block its intracellular trafficking and efflux activity. J Neurosci, 34, 14079–14095.

29. Moore, C., Sambrook, J., Walker, M., Tolkien, Z., Kaptoge, S., Allen, D., Mehenny, S., Mant, J., Angelantonio, E.D., Thompson, S.G. et al. (2014) The INTERVAL trial to determine whether intervals between blood donations can be safely and acceptably decreased to optimise blood supply: study protocol for a randomised controlled trial. Trials, 15, 363.

30. Staley, J.R., Blackshaw, J., Kamat, M.A., Ellis, S., Surendran, P., Sun, B.B., Paul, D.S., Freitag, D., Burgess, S., Danesh, J. et al. (2016) PhenoScanner: a database of human genotype-phenotype associations. Bioinformatics, 32, 3207–3209.

31. McLaren, W., Gil, L., Hunt, S.E., Riat, H.S., Ritchie, G.R., Thormann, A., Flicek, P. and Cunningham, F. (2016) The Ensembl Variant Effect Predictor. Genome Biol, 17, 122.

32. Lawrence, M., Huber, W., Pages, H., Aboyoun, P., Carlson, M., Gentleman, R., Morgan, M.T. and Carey, V.J. (2013) Software for computing and annotating genomic ranges. PLoS Comput Biol, 9, e1003118.

33. Yates, A., Akanni, W., Amode, M.R., Barrell, D., Billis, K., Carvalho-Silva, D., Cummins, C., Clapham, P., Fitzgerald, S., Gil, L. et al. (2016) Ensembl 2016. Nucleic Acids Res, 44, D710–D716.

34. Pare, G., Chasman, D.I., Parker, A.N., Zee, R.R., Malarstig, A., Seedorf, U., Collins, R., Watkins, H., Hamsten, A., Miletich, J.P. et al. (2009) Novel associations of CPS1, MUT, NOX4, and DPEP1 with plasma homocysteine in a healthy population: a genome-wide evaluation of 13 974 participants in the Women’s Genome Health Study. Circ Cardiovasc Genet, 2, 142–150.

35. van Meurs, J.B., Pare, G., Schwartz, S.M., Hazra, A., Tanaka, T., Vermeulen, S.H., Cotlarciuc, I., Yuan, X., Malarstig, A., Bandinelli, S. et al. (2013) Common genetic loci influencing plasma homocysteine concentrations and their effect on risk of coronary artery disease. Am J Clin Nutr, 98, 668–676.

36. Mudd, S.H., Uhlendorf, B.W., Freeman, J.M., Finkelstein, J.D. and Shih, V.E. (1972) Homocystinuria associated with decreased methylenetetrahydrofolate reductase activity. Biochem Biophys Res Commun, 46, 905–912.

37. Elmore, B.O., Bollinger, J.A. and Dooley, D.M. (2002) Human kidney diamine oxidase: heterologous expression, purification, and characterization. J Biol Inorg Chem, 7, 565–579.

38. Sanna, S., Busonero, F., Maschio, A., McArdle, P.F., Usala, G., Dei, M., Lai, S., Mulas, A., Piras, M.G., Perseu, L. et al. (2009) Common variants in the SLCO1B3 locus are associated with bilirubin levels and unconjugated hyperbilirubinemia. Hum Mol Genet, 18, 2711–2718.

39. Kaplan, M., Hammerman, C., Vreman, H.J., Stevenson, D.K. and Beutler, E. (2001) Acute hemolysis and severe neonatal hyperbilirubinemia in glucose-6-phosphate dehydrogenase-deficient heterozygotes. J Pediatr, 139, 137–140.

40. Yang, H., Robinson, P.N. and Wang, K. (2015) Phenolyzer: phenotype-based prioritization of candidate genes for human diseases. Nat Methods, 12, 841–843.

41. Cheng, D., Knox, C., Young, N., Stothard, P., Damaraju, S. and Wishart, D.S. (2008) PolySearch: a web-based text mining system for extracting relationships between human diseases, genes, mutations, drugs and metabolites. Nucleic Acids Res, 36, W399–405.

42. van Driel, M.A., Bruggeman, J., Vriend, G., Brunner, H.G. and Leunissen, J.A. (2006) A text-mining analysis of the human phenome. Eur J Hum Genet, 14, 535–542.

43. Hristovski, D., Peterlin, B., Mitchell, J.A. and Humphrey, S.M. (2005) Using literature-based discovery to identify disease candidate genes. Int J Med Inform, 74, 289–298.

44. Van Vooren, S., Thienpont, B., Menten, B., Speleman, F., De Moor, B., Vermeesch, J. and Moreau, Y. (2007) Mapping biomedical concepts onto the human genome by mining literature on chromosomal aberrations. Nucleic Acids Res, 35, 2533–2543.

45. Yu, W., Wulf, A., Liu, T., Khoury, M.J. and Gwinn, M. (2008) Gene Prospector: an evidence gateway for evaluating potential susceptibility genes and interacting risk factors for human diseases. BMC Bioinformatics, 9, 528.

46. Bornigen, D., Tranchevent, L.C., Bonachela-Capdevila, F., Devriendt, K., De Moor, B., De Causmaecker, P. and Moreau, Y. (2012) An unbiased evaluation of gene prioritization tools. Bioinformatics, 28, 3081–3088.

47. Chen, J., Bardes, E.E., Aronow, B.J. and Jegga, A.G. (2009) ToppGene Suite for gene list enrichment analysis and candidate gene prioritization. Nucleic Acids Res, 37, W305–311.

48. Maurano, M.T., Humbert, R., Rynes, E., Thurman, R.E., Haugen, E., Wang, H., Reynolds, A.P., Sandstrom, R., Qu, H., Brody, J. et al. (2012) Systematic localization of common disease-associated variation in regulatory DNA. Science, 337, 1190–1195.

49. Aerts, S., Lambrechts, D., Maity, S., Van Loo, P., Coessens, B., De Smet, F., Tranchevent, L.C., De Moor, B., Marynen, P., Hassan, B. et al. (2006) Gene prioritization through genomic data fusion. Nat Biotechnol, 24, 537–544.

